# Real-time tracking of intracellular prenyl phosphate pools in the marine diatom *Phaeodactylum tricornutum* with a metabolite protein-based biosensor

**DOI:** 10.1101/2025.03.31.646338

**Authors:** Payal Patwari, Tessa Moses, Michele Fabris

## Abstract

Metabolite-responsive, protein-based biosensors are a powerful tool for monitoring cellular metabolite dynamics *in vivo* and accelerating strain engineering workflows in microorganisms. In this study, we introduced a previously developed protein-based biosensor, computationally designed to detect farnesyl diphosphate (FPP), in the marine diatom *Phaeodactylum tricornutum.* We expressed two versions of the biosensor constitutively, under a strong promoter-terminator pair using extrachromosomal episomes, and we parameterized the capacity of both designs in detecting intracellular metabolite levels. Initial assays revealed that the two versions of the biosensor we investigated, S3-2D and S3-3A, had specificity not only for FPP but also for other exogenously supplied prenyl phosphates such as geranyl diphosphate (GPP) and geranylgeranyl diphosphate (GGPP) in a dose-dependent manner. We further demonstrated the capacity of S3-3A to track perturbations in the endogenous prenyl phosphate pools by testing it in the presence of pharmacological inhibition of the mevalonate pathway. Moreover, S3-3A generated signal “*hot-spots*” around the peroxisomes, suggesting their involvement in isoprenoid biosynthesis, which led us to characterize the subcellular localization of the key enzyme mevalonate kinase. These findings lay the groundwork for developing metabolite-responsive biosensors as robust tools for monitoring and investigating prenyl phosphate dynamics, providing a foundation for advanced metabolic engineering of microalgae.

## Introduction

A biosensor is an analytical device that combines a biological component with a detector or regulator to measure the presence or concentration of specific metabolites, enzymes, or nucleic acids, *in vitro* or *in vivo*^1^. Protein-based biosensors function by binding to a ligand, which triggers a reporter (for example bioluminescence or fluorescent proteins) to produce a measurable signal, which in turn serves as a proxy for the presence of the target molecule^2^. Protein-based biosensors have drastically accelerated the investigation of fundamental traits and facilitated metabolic engineering in established model microorganisms such as *Escherichia coli* and *Saccharomyces cerevisiae*, with applications in quantifying and monitoring metabolites in a concentration-dependent manner^3^, high-throughput screening ^4^, and elucidating metabolic dynamics and pathway regulation^5,6^. However, the development and implementation of such biosensors in microalgae has never been reported.

While eukaryotic microalgae are becoming attractive hosts for photosynthesis-driven bioproduction through synthetic biology and metabolic engineering, their genetic and biological complexity, as well as the still limited genetic toolbox, often limit the reach of metabolic engineering. Microalgal metabolic pathways and regulatory networks are often compartmentalized and different than those of more established hosts, and are also less characterized, requiring comprehensive knowledge of their location and function to enable effective engineering strategies^7,8^. Despite these challenges, increasing efforts are being invested to establish molecular tools for microalgae to redesign metabolic pathways to enhance productivity and efficiency, especially in marine diatoms, which possess attractive biological traits and have advanced genetic tractability^9,10^.

Diatoms are heterokont microalgae with complex and diverse metabolism, and can be genetically engineered to produce heterologous compounds, including terpenoids^11,12^. These find application as pharmaceuticals, fragrance and flavor ingredients, cosmetics and biofuels^13^. Terpenoids are derived from the prenyl phosphates geranyl diphosphate (GPP), farnesyl diphosphate (FPP), and geranylgeranyl diphosphate (GGPP), via the mevalonate pathway (MVA) and the methyl-erythritol phosphate (MEP) pathway. In *Phaeodactylum tricornutum*, similar to plants and different from bacteria and yeasts, both MVA and MEP operate to fuel the biosynthesis of pigments such as sterols and carotenoids, respectively^14^. In some species of *Pseudonitzschia*, isoprenoids such as geranyl diphosphate (GPP) have a relevant ecological and environmental role, as they are precursors to secondary metabolites such as the toxin domoic acid, at the basis of harmful algal blooms (HABs)^15^. The organization of these pathways have been mapped with genomic^9,16^ and transcriptomic data^17^, highlighting the presence and function of novel enzymes^16,18^, however, their regulation and detailed subcellular localization is still largely unknown.

Recent advancements in genetic engineering tools for diatoms include resources such as extrachromosomal episomal expression^19^, which offers a relatively stable and rapid platform to develop and prototype complex synthetic genetic functions through rapid iterations of *Design-Build-Test-Learn* (DBTL) cycles. This is further accelerated by the development of modular DNA assembly workflows^20,21^ and high-throughput screening approaches^21–24^.

Profiling of engineered algal strains and metabolite analyses are commonly carried out using gas or liquid chromatography coupled to mass spectrometry (GC-MS or LC-MS), which require full-scale experiments, are costly, resource-intensive, and require destructive extraction methods. This ultimately hampers engineering workflows by slowing down the DBTL cycles. Previous research has demonstrated the feasibility of engineering plant transgenes into diatoms for the successful production of monoterpenoids^9,21^, sesquiterpenoids^25^, triterpenoids^12^, carotenoids^26^, and cannabinoids^10^. However, the yields have been relatively low and limited, likely due to limited precursor availability and metabolic flux imbalance, among other factors.

Biosensors can overcome some of these bottlenecks, reducing the need for conventional assays to identify key metabolic elements and fluxes, thus increasing the speed and efficiency of strain engineering^27,28^. Metabolite biosensors are compatible with the genetic toolbox of diatoms to enable rapid, non-invasive detection of metabolites, facilitating high-throughput phenotyping and the investigation of the intracellular environment, monitoring single-cell productivity^2^ and understanding metabolite dynamics. Integration of flow cytometry, fluorescent activated cell sorting (FACS) and microtiter plate reader assays further supports the miniaturization and acceleration of experimental workflows.

In this work, we introduced a computationally designed metabolite-responsive protein-based biosensor^29^ in *P. tricornutum*. Ankyrin repeat (AR) containing proteins can mediate heterodimeric protein-protein interactions^30^. This function was harnessed by Glasgow and colleagues to build *de novo* binding sites into heterodimeric protein-protein interface of maltose binding protein (MBP) and AR protein to sense an important terpenoid precursor FPP. In their work, Glasgow and colleagues linked their best-performing protein pairs S3-2D and S3-3A, to a dimerization-dependent split fluorescent protein pair (ddGFP and ddRFP) to test the input-output response by *in vitro* expression^29^. Building on these resources, we tested both biosensors in *P. tricornutum* for real-time monitoring of exogenous and endogenous FPP and other structurally similar intermediates (GPP and GGPP). Furthermore, we demonstrated the utility of the biosensors to track subcellular metabolite *hotspots*, revealing key new elements in isoprenoid biosynthesis pathways of diatoms.

## Results and discussion

### The S3-2D and S3-3A protein biosensors respond to GPP besides FPP *in vitro*

From those developed by Glasgow et al., 2019, we chose the two biosensor designs which displayed a robust signal response to FPP: S3-2D (ddRFPb-AR-2.7 and MBP-2.5-ddGFPa) and S3-3A (ddRFPb-AR-2.7 and MBP-3.6-ddGFPa). We also employed WT (wild-type - ddRFPb-WTAR and WTMBP-ddGFPa), in which dimerization is not affected by the presence of FPP^29^ as control (Fig. 1). The biosensor labels are composed of scaffold (S3), design generation (2/3) followed by a letter (A/D)^29^. Before testing the biosensors’ functionality in diatoms, we assessed it *in vitro* in the presence of increasing concentrations (0 to 10 µM) of FPP, confirming the activity observed by Glasgow et al. (2019), and the suitability of our experimental set-up (Fig. S1A). Glasgow et al., (2019) demonstrated the response of the engineered biosensors to FPP; we hypothesized that the S3-2D and S3-3A biosensor variants could also respond to other prenyl phosphates since this aspect was not investigated by Glasgow and co-workers. Using the same experimental set-up, we profiled the response of both biosensor designs to increasing concentrations of GPP, structurally similar to FPP (Fig. S1B). S3-3A demonstrated broad-range sensitivity to FPP (0.1-1 µM) and GPP (0.1-2 µM), with a saturation trend at concentrations higher than 4 µM, whereas S3-2D showed sensitivity to FPP and GPP within 0.1-0.5 µM. S3-3A exhibited a wide dynamic range of sensitivity for both FPP and GPP and was reported as the best biosensor design by Glasgow et al., 2019 (Fig. S1).

**Figure 1.**
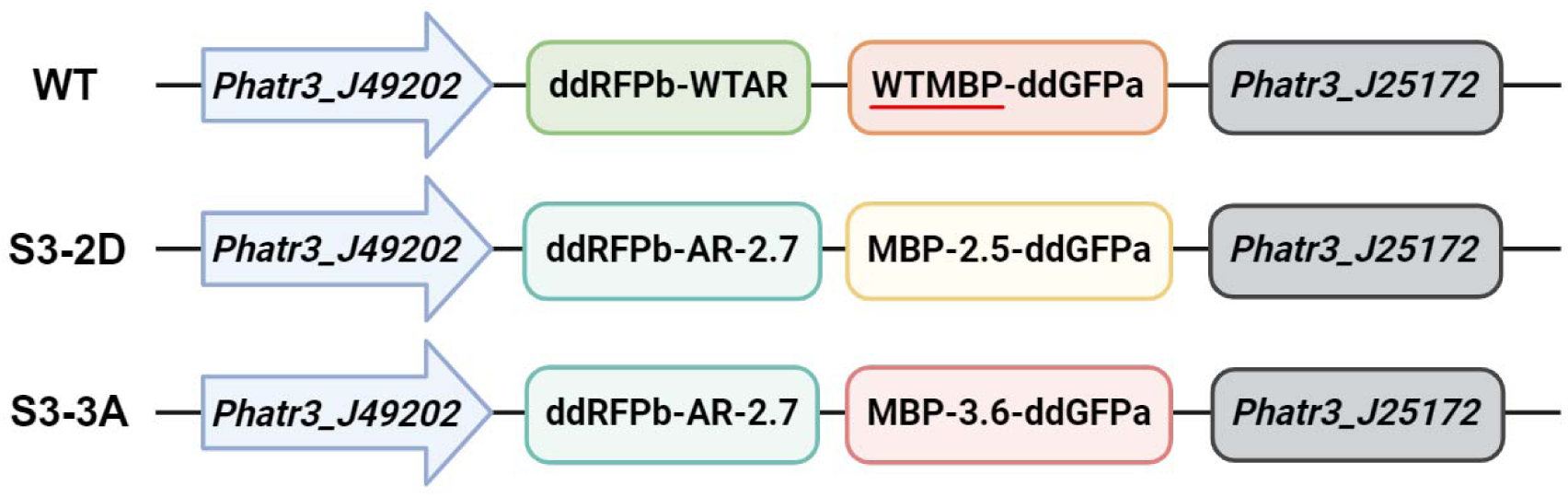
Schematic representation of construct design of WT, S3-2D and S3-3A (Glasgow et al., 2019) expressed as episomes in *P. tricornutum*. The biosensors were flanked by *Phatr3_J49202* and *Phatr3_J25172 (FcBPt)* promoter (blue) and terminator (grey) regions, respectively. Detailed mechanisms of action are described in (Glasgow et al., 2019).

### Extrachromosomal expression of S3-2D and S3-3A biosensors in diatoms

To test their functionality in diatoms, we expressed the dimeric biosensors S3-2D and S3-3A and WT (control) in *P. tricornutum* from extrachromosomal episomes as independent transcriptional units without targeting peptides (Fig. 1). To ensure strong and constitutive expression, both biosensor components were placed under the control of the promoter region of *Phatr3_J49202*^20^, in combination with the terminator region of *Phatr3_J25172* (*FcBPt*) (Fig. 1). This combination has previously been shown to ensure high and consistent expression levels^9^. We screened 30 independent transformant cell lines by flow cytometry, based on GFP fluorescence intensity and chlorophyll autofluorescence (Fig. S2). The GFP signal for S3-2D and S3-3A plausibly reflected a basal biosensor activity in response to endogenous cytosolic metabolite availability and thus preliminary evidence of the correct assembly of the protein complex. Among the 30 cell lines associated with detectable GFP signal, we selected three for each construct (S3-2D, S3-3A and WT). We profiled these in the late exponential growth phase (on day 5 of batch cultivation), and we ranked them based on mean GFP fluorescence intensity (MFI) (Fig. 2A). The signal response across cell lines expressing the same biosensor variant was relatively consistent, except for S3-3A-3, which showed a substantially lower signal than S3-3A-1 and S3-3A-2. Overall, our observations are aligned with previous expression performances of the extrachromosomal expression of recombinant proteins in the cytosol, flanked by the *Phatr3_J49202* promoter and the *Phatr3_J25172* terminator region pair^9^. Conversely, cell lines expressing S3-2D showed a marked difference in the measured signal, with generally lower MFI. The mean fluorescence intensities of both S3-2D and S3-3A reflect a combination of factors: the different expression levels of the biosensor genes and the endogenous pool of prenyl phosphates available in the cytosol. Assuming that the promoter-terminator pair ensures relatively similar expression levels and that diatom cultivated in the same conditions have similar cytosolic pools of prenyl phosphates and allow a degree of phenotypic variation in the population both in terms of expression level and metabolite accumulation (Fig. 2B), these preliminary observations suggested that the biosensor design S3-3A might perform better in the cytosolic environment of *P. tricornutum*, even though the two biosensor variants showed very similar performances *in vitro* (Fig. S1) and in Glasgow et al., 2019. The flow-cytometry analyses of selected cell lines expressing either S3-2D, S3-3A or WT provided a useful initial assessment of the behavior of the different biosensors in the diatom cytoplasm. S3-3A exhibited the highest GFP fluorescence intensity compared to S3-2D (Fig. 2B) and WT. These intensities were consistently observed across the three independent lines tested, except for S3-2D-1 and S3-2D-3, where the intensities were like WT. These results aligned with the report of Glasgow et al. (2019), where S3-3A was an improved version of S3-2D. Specifically, S3-3A was designed with Y197A mutation in the MBP component that stabilized the ternary complex and was reported to be the best biosensor variant to detect FPP at nanomolar levels^29^. Based on the screening, the cell lines S3-2D-2, S3-3A-1, and S3-3A-2 with high MFI (Fig. 2A) were selected for further characterization, with WT as a control.

**Figure 2.**
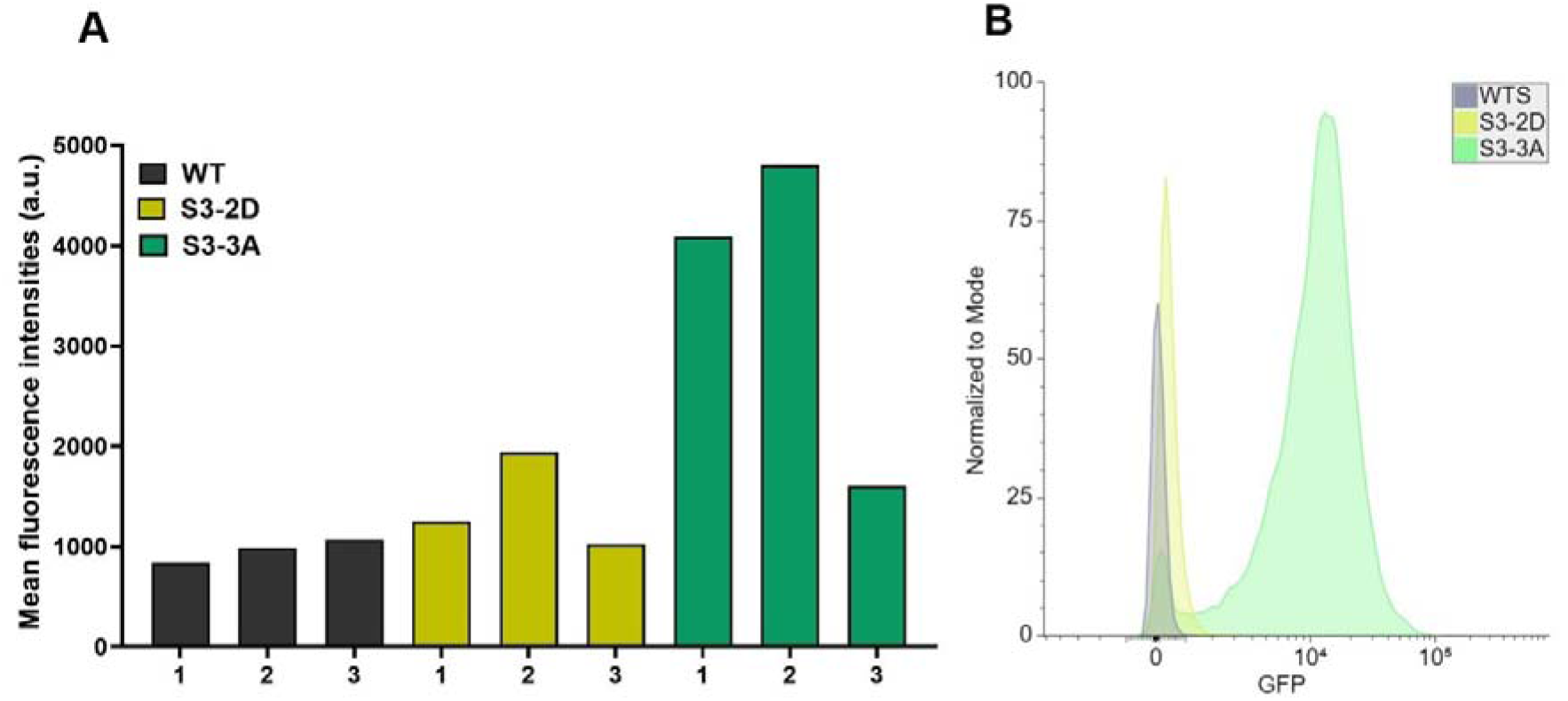
(A) Flow cytometry-based screening of transgenic diatom cell lines expressing WT, S3-3A or S3-2D biosensors. The bars represent the mean fluorescence intensity of 10,000 cells for each sample, numbers on the x-axis indicate independent cell lines. (B) Representative histograms of one clonal transformant cell line expressing WT, S3-3A or S3-2D biosensors, showing flow cytometry populations of cells based on GFP fluorescence intensity. Cell populations of WT and S3-2D and S3-3A were normalized to the mode (n=10,000).

### Intracellular S3-2D and S3-3A biosensors respond to different prenyl phosphates in diatoms

To evaluate the functionality, sensitivity, operational range, and detection specificity of the biosensors in the diatom intracellular environment, we subjected cell lines S3-2D-2, S3-3A-1, S3-3A-2, and WT to increasing macro-concentrations (0-1000 µM) of GPP, FPP, as well as GGPP in 96-well plates over 24 and 48 hours (Fig. 3). While GGPP was not previously tested *in vitro* in our previous assays, we included it in this set of experiments, hypothesizing that S3-2D and S3-3A could react upon binding to it, based on our previous results using GPP (Fig. S1). The fluorescence was measured both spectrophotometrically and by flow cytometry 3, 6, 24 and 48 hours after the addition of prenyl phosphates. In this experiment, we observed the capacity of *P. tricornutum* cells to incorporate extracellular prenyl phosphates besides GPP, whose uptake was recently demonstrated^10^.

**Figure 3.**
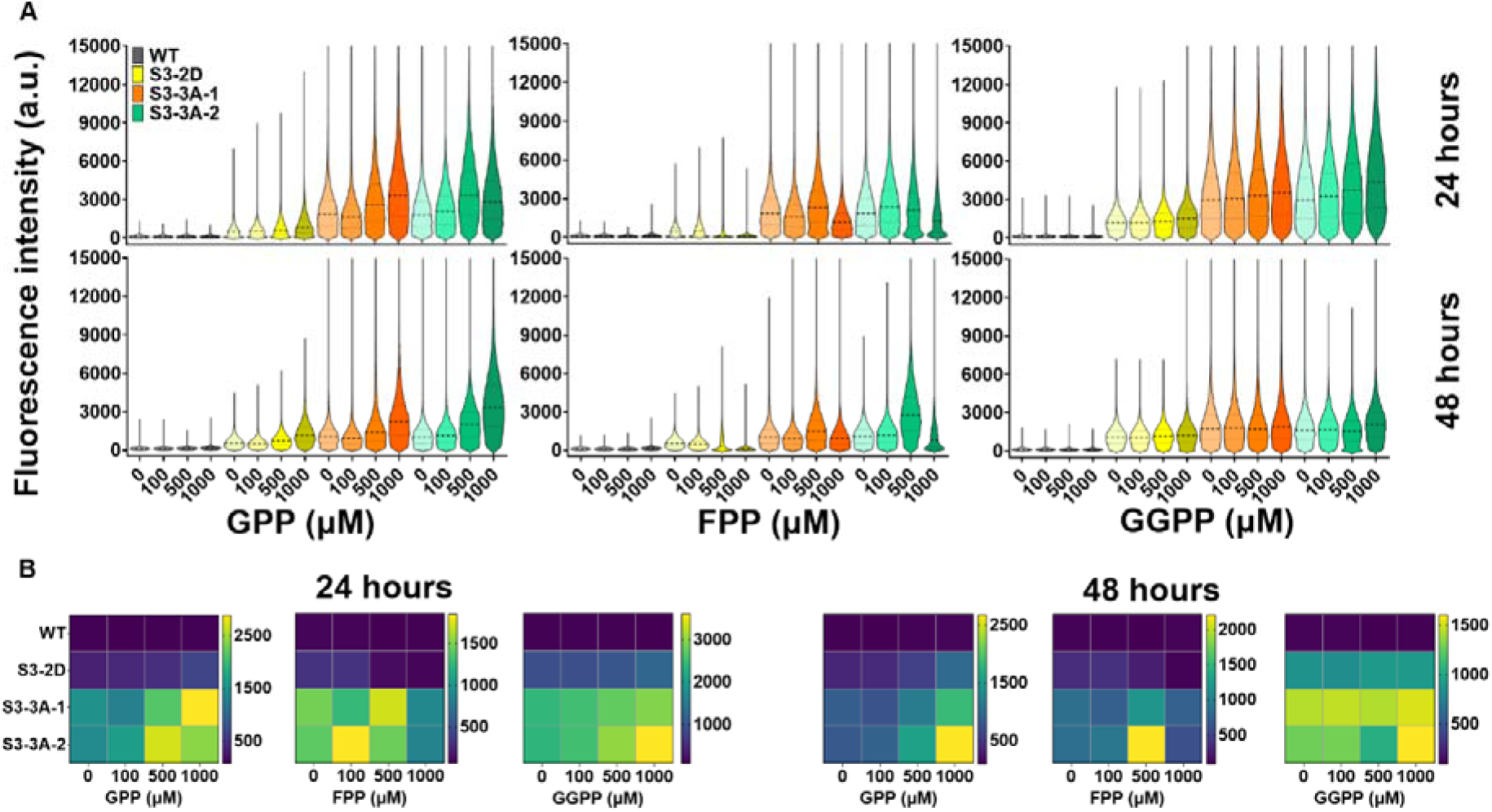
(A) Violin plots depicting the activity of biosensor variants S3-2D and S3-2D, and WT after 24 hours and 48 hours of incubation after the addition of macro-concentrations of GPP, FPP and GGPP (n=30,000). Dashed black lines represent the median and grey dotted lines represent the upper and lower quartiles. (B) Heatmaps depicting the geometric mean of the fluorescence intensities after 24 and 48 hours of incubation

Prior to the addition of prenyl phosphates, we observed that all cell lines expressing S3-2D or S3-3A showed a basal level of signal presumably due to the endogenous prenyl phosphate pools (Fig. 2B). WT showed a lower, basal GFP signal due to the natural lower affinity of AR and MBP proteins, independently from the presence of prenyl phosphates^29^. The GFP fluorescence intensity was measured by flow cytometry for all diatom cell lines expressing WT, S3-2D and S3-3A after 3, 6, 24, 48 hours of incubation. After 3 (data not shown) and 6 hours of incubation with different concentrations of GPP, FPP and GGPP, we could not detect changes in the fluorescence intensity of WT, S3-2D and S3-3A (Fig. S3).

After 24 hours of incubation with exogenous GPP, FPP and GGPP, we observed a dose-response increase in fluorescence intensity at 100 µM in the presence of FPP, and from 100-1000 µM for GPP and GGPP in diatoms expressing S3-2D or S3-3A (Fig. 3A and B). Without the addition of any metabolite, S3-2D exhibited significantly lower fluorescence signal intensity than S3-3A, associated with a defined distribution around the median, indicating a rather homogeneous phenotypic population in response plausibly to endogenous metabolite levels. In contrast, both cell lines expressing S3-3A showed markedly higher fluorescence intensity, associated with a more heterogeneous population in the same conditions (Fig. 3A). After 48 hours, we observed an overall decrease in the fluorescence intensity of S3-2D, S3-3A-1 and S3-3A-2 compared to 24 hours, except when GPP was added. In this case, we observed an increase in median and geometric mean fluorescence for S3-3A-1 and S3-3A-2 lines (Fig. 3A and B). After 48 hours of incubation with prenyl phosphates, all lines exhibited more defined and homogenous populations in relation to their output signal phenotype, except for S3-3A-2 which showed broad distribution at higher concentrations of 500-1000 µM of GPP and 500 µM of FPP (Fig. 3A). We observed that FPP at a concentration of 500 µM led to a substantial decrease in the biosensor signal after 48 hours, and already after 24 hours at 100 µM, presumably due to the toxic effects of FPP on diatom physiology. While this phenomenon has never been observed in diatoms before, this aligns with previous reports indicating that FPP is toxic to yeast and bacteria at concentrations above 200 µM^31–34^.

The kinetics of the output of biosensors and the stability over time is a key parameter and depends on the design and functioning of the biosensor itself, its interaction with the ligand, and the cellular environment of the host organism. From our observations, the performances of the S3-2D and S3-3A upon the addition of exogenous ligands in *P. tricornutum* are comparable with those of previously reported biosensors with other hosts^35,36^. For instance, the fluorescence intensity of a plasmid-based biosensor to detect exogenously supplied glucuronate in *E. coli* was highest at 10 hours and levelled off between 12 and 16 hours of cultivation^37^. In another study, a yeast transcription factor-based biosensor was stably integrated into mammalian cells and reported a four-fold increase in fluorescence within 4 hours of extracellular ligand feeding which rose exponentially within 24-48 hours^38^.

The dynamic range, defined as the concentration window between the minimum and maximum biosensor output compared to the untreated fluorescence intensity^39^, was in the range of 50-500 µM FPP for S3-3A-2 and 100-500 µM GPP and GGPP for S3-2D, S3-3A-1 and S3-3A-2 (Fig. 3A and 4). Our experiments demonstrated that both S3-2D and S3-3A protein biosensors, while computationally designed to bind to FPP, are capable of binding other structurally similar prenyl phosphates GPP and GGPP. Hence, both S3-2D and S3-3A biosensors offer precise proxies to detect total pools of prenyl phosphates in *P. tricornutum* cells. The fluorescence intensity of both biosensors exhibited a dose-dependent response with increasing concentrations of GPP, even after 48 hours of incubation. However, the intensity decreased for both lines after 48 hours with FPP and GGPP (Fig. 3A and B). This observation suggests a cumulative effect, possibly due to the direct binding of the biosensors to GPP or FPP, if this is formed in two steps from GPP, as in *S. cerevisiae*, condensing with a molecule of IPP^16^. In *P. tricornutum*, the exact mechanism governing the biosynthesis of prenyl phosphates is still elusive, albeit candidate genes have been identified and based on this we could hypothesize such a mechanism^9^. In this experiment, the S3-3A design exhibited a more pronounced signal intensity increase in response to 100 µM FPP, and 1000 µM GPP and GGPP compared to S3-2D. The latter, despite having similar activity to S3-3A *in vitro*, clearly shows a different behavior *in vivo*, when expressed in diatoms. The intensity of the signal was markedly lower than S3-3A, while still showing a correlation between metabolite concentration and signal intensity, with better performances in response to GGPP, followed by GPP and finally FPP, suggesting that S3-2D might have a higher specificity for GGPP (Fig. 3A). Based on our results, we cannot exclude the possibility that S3-3A and S3-2D may respond also to other metabolites, which are structurally different but related to prenyl phosphates.

**Figure 4.**
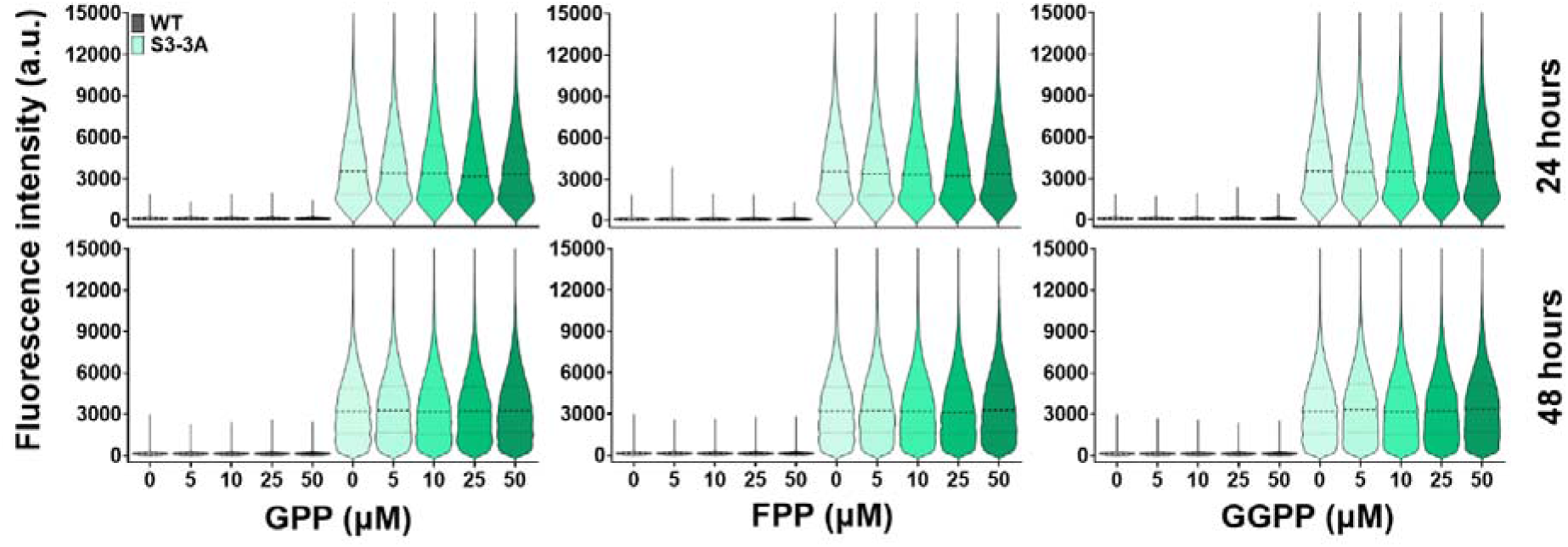
Violin plots depicting the activity of WT and S3-3A after 24 hours and 48 hours of incubation after the addition of micro-concentrations of GPP, FPP and GGPP (n=30,000). Dashed black lines represent median and grey dotted lines represent upper and lower quartiles.

To provide a complete, *in-depth* parameterization of the biosensors in diatoms, we analyzed their performance at the single-cell level in the clonal population, which is enabled by flow cytometry (Fig. 3A). This resolution highlighted substantially heterogeneous populations within the same clonal cell lines, which included cells that did not emit signals as well as cells that emitted very high levels of fluorescence. This effect was more prominent in samples expressing S3-3A, while S3-2D expressing cell lines were characterized by a more defined population. Overall, the comparison of the median fluorescence values of each clonal cell line population (Fig. 3A) showed a mean response proportional to the prenyl phosphate concentration. While the analysis of diatoms expressing the biosensors with flow cytometry allows to profile the response with a single-cell resolution and to appreciate the difference in the population homogeneity, the geometric mean of the fluorescent signal of each cell in a sample can be used as a simpler, but still valid proxy to convey information on the different levels of metabolites (Fig. 3B). This could resemble the typically “averaged” output of measurement methods that are less sophisticated and more accessible than flow cytometry, such as those deriving from spectro/photo/fluorometers, possibly simplifying and accelerating screening and strain improvement workflows. To demonstrate this, we profiled the biosensors’ output using a microplate spectrofluorometer, equipped for GFP fluorescence detection. While the microplate reader only provides an estimate of average fluorescence intensity of the entire population, it offers robustness and accessibility, particularly in settings lacking high-end instruments like flow cytometry^40^. The geometric mean fluorescence intensities acquired from flow cytometry after the addition of a range of prenyl phosphates corresponded to the values obtained as normalized fluorescence/OD_750_ by a microplate reader (Fig. S4).

The sensitivity of a biosensor is its ability to effectively detect small changes in metabolite concentrations and the operational range is the significant change in the biosensor output defining the lower and upper detection limits^41^. For biosensors to be useful for screening metabolites *in vivo*, these should be highly sensitive, and the operational range should be within the limits of reported physiologically relevant concentrations of the host. Since microalgae^9,42,43^ and other industrially relevant microbial chasses^34^ are reported to have low levels of endogenous isoprenoid precursors, in the order of nanomolar concentrations^44^, we investigated the sensitivity of the selected S3-3A at lower concentrations of prenyl phosphates (0-50 µM)^45^. 24 hours after the addition of exogenous prenyl phosphates, we could not observe any deviation from the basal fluorescent signal, in response to the endogenous pool of metabolites that could be related to the addition of exogenous prenyl phosphates (Fig. 4). Similarly, after 48 hours we could not observe differences in mean fluorescence in the response between samples to which prenyl phosphates were added and the controls. However, after 48 hours, biosensor-expressing diatoms assumed a wider population spread compared to the distributions observed after 24 hours of incubation, presumably reflecting differences in the intracellular prenyl phosphate pool.

The detection limit for S3-3A in the cytosol of *P. tricornutum* was 50 µM of exogenously added metabolites, indicating that its sensitivity range is low. However, these results are directly affected by the stability and availability of prenyl phosphates in artificial seawater, and by the yet unknown uptake rate of diatoms. This might depend on the presence, activity, and affinity of putative transporters and plausibly low at micro-concentrations. Since intracellular prenyl phosphate pools have been reported at low concentrations in wild-type *P. tricornutum* cells in the range of 0.2-0.5 µg/million cells^9^ and 0.1-0.4 µg/g dry cell weight in *S. cerevisiae*^46^, further work is needed to fine-tune the biosensor’s performance to maximize the sensitivity. Overall, from the above results the operational range of S3-3A was observed as 50-500 µM for FPP and GGPP (Fig. 3B and 4) and 500-1000 µM for GPP (Fig. 3B). Notably, we observed a consistent difference in the mean fluorescence intensity between transgenic diatom strains expressing WT and S3-2D (about 2-fold), and between WT and S3-3A (5-fold) (Fig. 3A and 5A), compared to the initial screening experiments (Fig. 2A), where the difference was 1-fold and 4-fold, respectively. These latter experiments were carried out 14 months after the screening and selection, and diatom strains expressing the WT dimeric complex might have lost the capacity to express the WT construct, as confirmed in later experiments (Fig. 6D). Conversely, the basal level of signal exhibited by diatom lines expressing either S3-2D and S3-3A constructs remained consistent throughout the series of experiments. Due to the overall superior signal intensity response, we selected the biosensor S3-3A for all further experiments. We focused on S3-3A-2 as a representative cell line, henceforth referred to as S3-3A.

**Figure 5:**
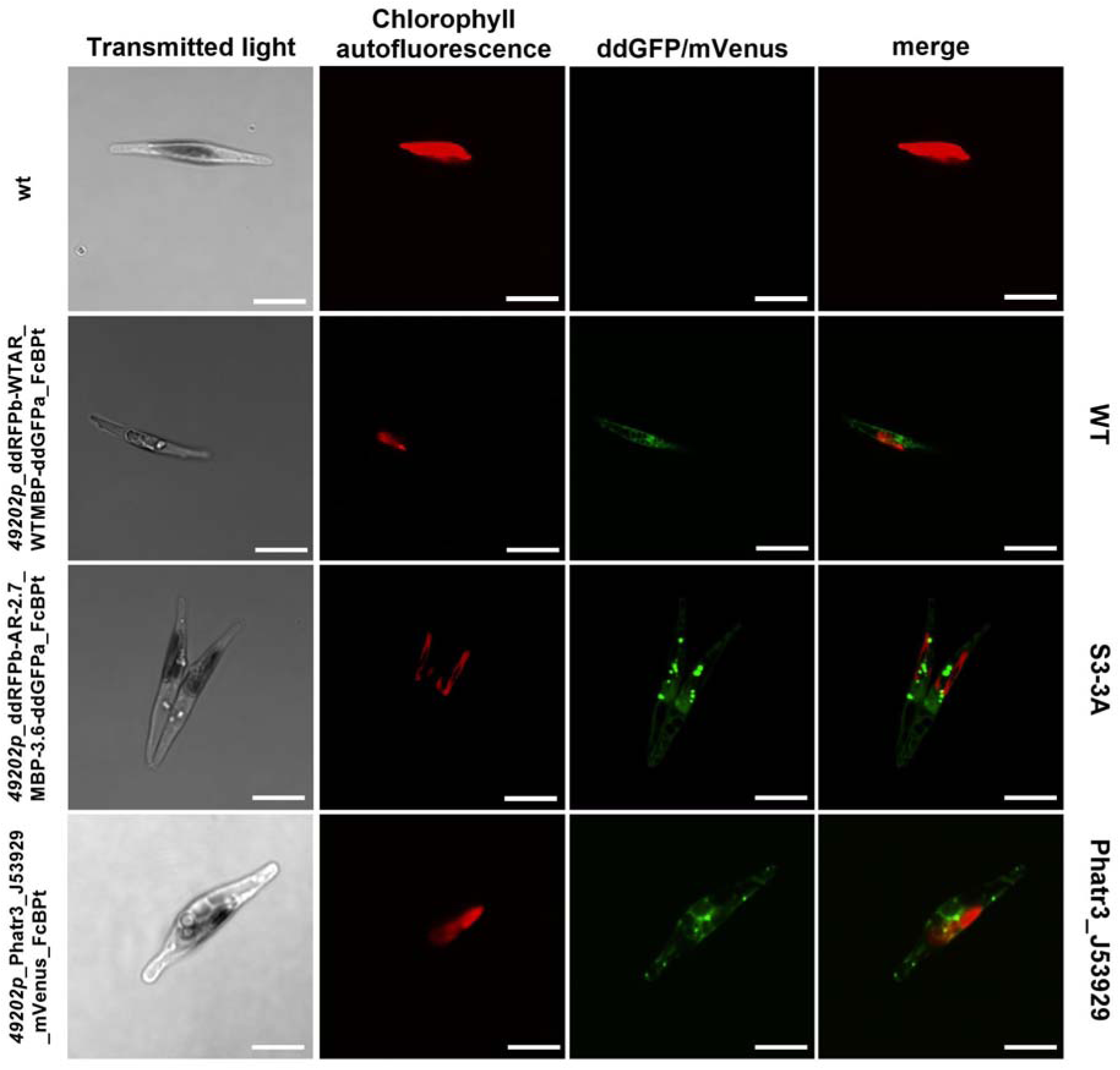
Subcellular localization of ddRFPb-WTAR_WTMBP-ddGFPa (WT), ddRFPb-AR-2.7_MBP-3.6-ddGFPa (S3-3A) and Phatr3_J53929_mVenus (PtMVK) after four days of cultivation; scale bars=5 µm. Images are representative of multiple observations.

**Figure 6.**
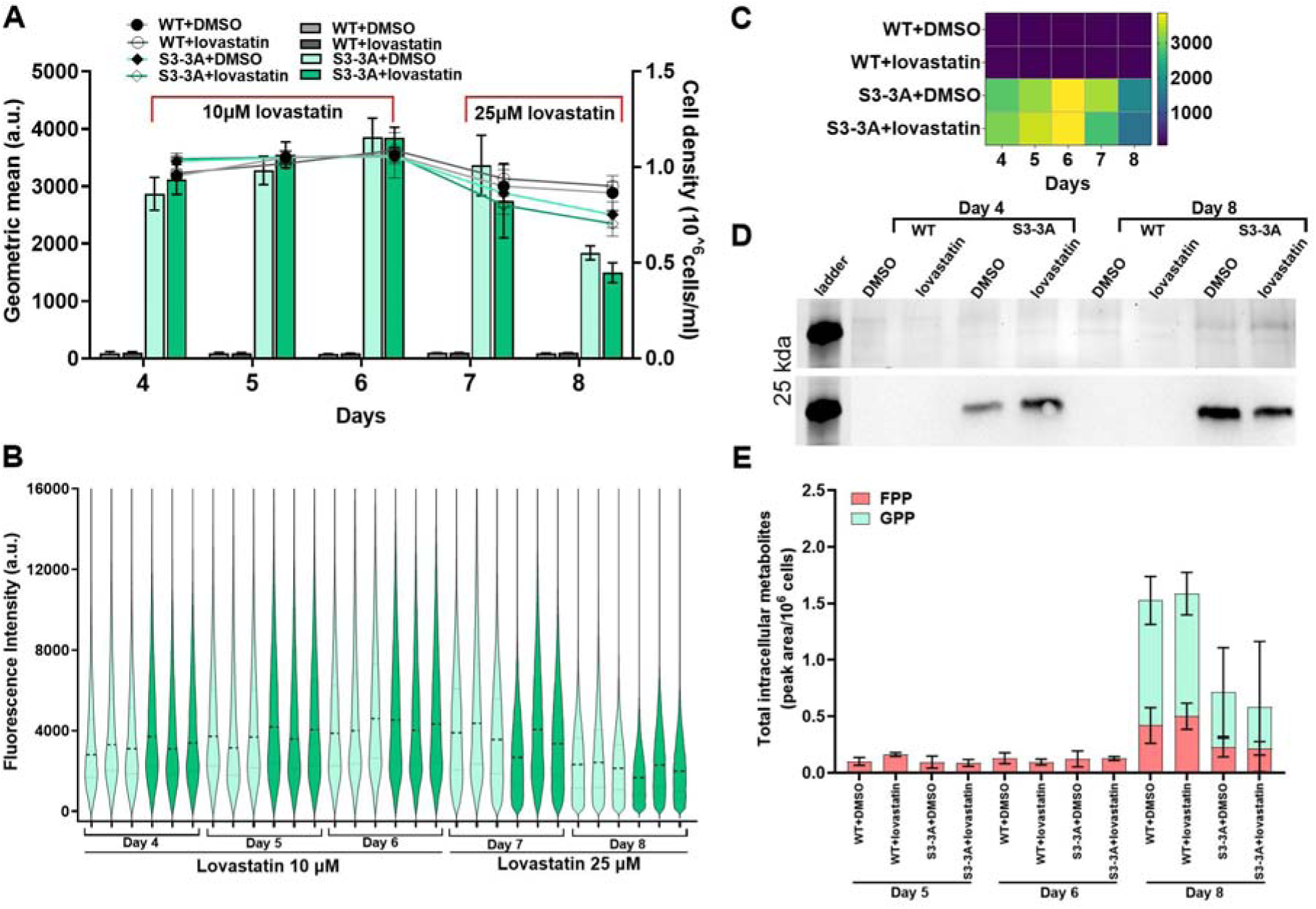
(A) Growth curves of WT and S3-3A strains plotted against cell density and geometric mean over 8 days with and without lovastatin (n=3). (B) Violin plot showing the fluorescence intensities over day 4 to day 8 within the GFP-gated population for S3-3A. Dashed black lines represent the median and grey dotted lines represent the upper and lower quartiles (n=30000 events). (C) Heatmap depicting the mean fluorescence intensity of WT and S3-3A with and without inhibition from day 4 to day 8 (n=3). (D) Immunoblot detection of the recombinant ddGFPa components with anti-GFP antibodies on pooled soluble protein fractions (n=3) collected on day 4 and day 8 in the presence or absence of lovastatin 25 µM. The upper panel reports an SDS-PAGE of the total soluble protein fraction as loading control; the lower panel shows the abundance of WT and S3-3A on day 4 and day 8. The expected size of ddGFPa component is 26 kDa. (E) Relative abundance of total intracellular metabolites (GPP, FPP and GGPP) analyzed by LC-MS in WT and S3-3A cells on days 5, 6 and 8 with either presence or absence of lovastatin.

### S3-3A effectively tracks prenyl phosphate biosynthesis and trafficking, and reveals the involvement of peroxisomes in the mevalonate pathway

The S3-3A and WT constructs lacked targeting peptide sequences, and their accumulation was expected to be cytosolic. Confocal fluorescence microscopy confirmed this subcellular localization through a diffused cytosolic fluorescence, revealing the formation of hotspots in cells expressing S3-3A, unlike those expressing WT (Fig. 5). These hotspots, marked by increased GFP signal, appeared as numerous small, spherical, and mobile structures within the cytosol. We hypothesize that they represent regions of elevated prenyl phosphate concentration and that the variably localized spherical structures observed in S3-3A-expressing cells correspond to peroxisomes ^54,55^. Peroxisomes have been shown to be involved in the MVA pathway in mammals^56–58^ and plants^59–62^. The acetyl CoA pools from β-oxidation in yeast peroxisomes have been exploited for terpenoid engineering due to peroxisomes’ proximity to the ER^63–65^. In diatoms, the localization of MVA pathway enzymes is not characterized and is generally considered to occur between the cytosol and ER. None of the putative prenyl transferases directly involved in the biosynthesis of GPP, FPP, and GGPP (Phatr3_J47271, Phatr3_J49325, Phatr3_J19000, Phatr3_J15180, Phatr3_J16615) are predicted to localize in peroxisomes. However, we manually identified a putative peroxisomal targeting sequence 2 (PTS2) in the enzyme mevalonate kinase (PtMVK, Phatr3_J53929, Fig. S7) based on the nonapeptide consensus [RKSH]-[LVIQAT]-X_5_-[HDQ]-[LAF] inferred from several known PTS2 peptides from model organisms^66,67^. When expressed in diatoms fused to mVenus at its C-terminal, we confirmed the localization of PtMVK to both the cytosol and peroxisomes, with a pattern similar to that observed in cells expressing S3-3A (Fig. 5). This indicates that peroxisomes are involved in isoprenoid biosynthesis, specifically in the MVA pathway. The S3-3A signal hotspots we observed might correspond to areas in the cytosol with higher prenyl phosphate concentrations, such as around the peroxisomes. Interestingly, the PtMVK enzyme contains a PTS2 motif, previously reported not to be recognized in *P. tricornutum*^68^.

### The S3-3A biosensor responds to variations of the cytosolic pool of prenyl phosphates in diatoms and to pathway inhibition

Next, we tested S3-3A for real-time tracking of the cytosolic endogenous prenyl phosphate pools and its capacity to detect their fluctuations *in vivo*. We investigated the response of S3-3A on endogenous prenyl phosphates either in the presence or absence of pharmacological inhibition of the MVA pathway, which prevalently contributes to the cytosolic GPP^9^ and FPP pools^16,69,70^ in *P. tricornutum*. The enzyme 3-hydroxy-3-methyl-glutaryl-CoA reductase (HMGR, EC:1.1.1.34) catalyzes the conversion of HMG-CoA to mevalonate, a rate-limiting step in the mevalonate pathway^16,17^. *P. tricornutum* possesses an HMGR enzyme, encoded by the gene *Phatr3_J16870*^71^ as part of the MVA pathway^16^ that provides FPP for the biosynthesis of sterols, essential components of the cytoplasmic membrane. In most MVA pathway harboring organisms such as fungi^72^, plants^73^, oomycetes^74^ and diatoms^69^, the primary effect of HMGR inhibition is generally reflected in decreased growth performances and lowered sterol levels. To perturb the isoprenoid MVA pathway, we treated diatom cultures with increasing concentrations of lovastatin (10 to 25 µM) and tracked the size of the cytosolic prenyl phosphate pools with S3-3A over 8 days of cultivation through the biosensor signal (Fig. 6A, B and C) and compared with the quantification of extracted metabolites by LC-MS (Fig. 6E) from day 5 to day 8. Additionally, we assessed the effect of the metabolic perturbation on the accumulation of pigments and sterols (Fig. 7), and to ensure that the biosensor signal predominantly reflected differences in metabolite availability, we monitored the expression of the ddGFPa recombinant constructs with immunoblots (Fig. 6D).

**Figure 7:**
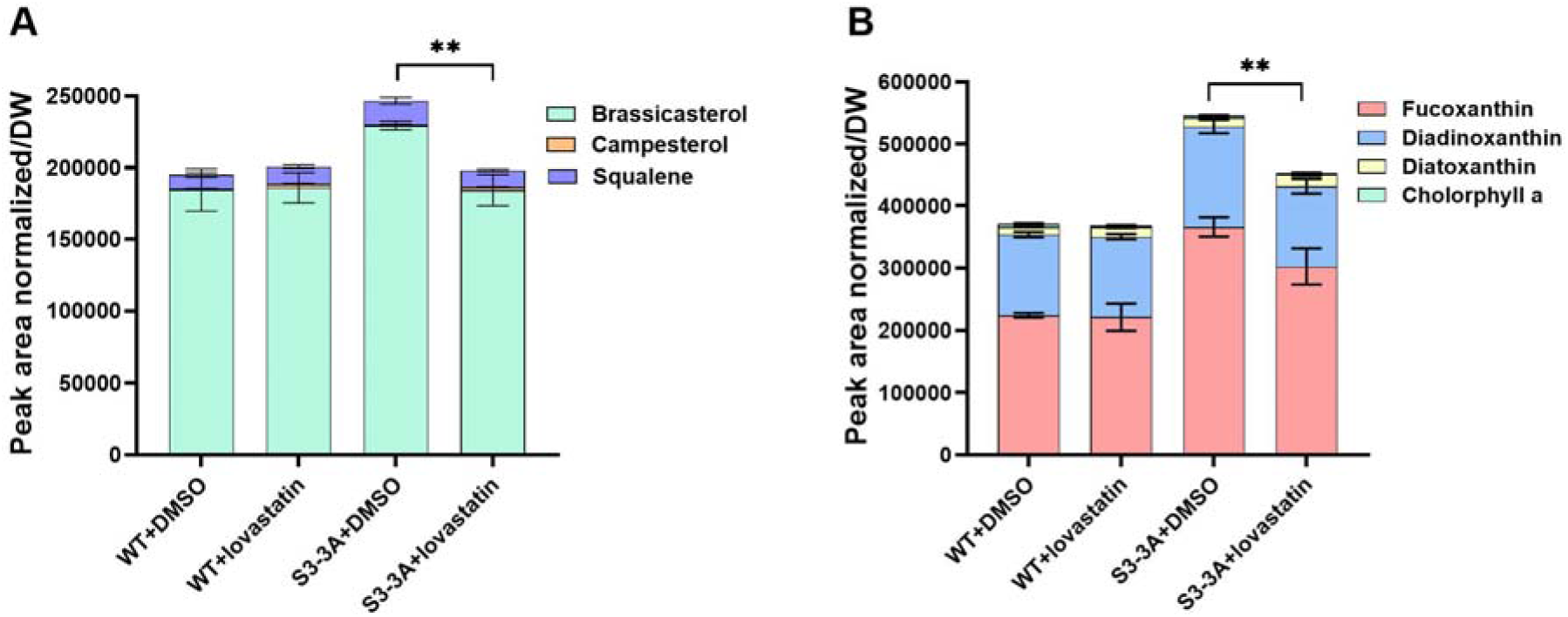
(A) Squalene, main sterols and (B) main carotenoids in diatoms expressing WT or S3-3A, measured at day 8 to demonstrate effective HMGR inhibition in S3-3A expressing cultures (n=3, one-way ANOVA test on total carotenoids and sterols, P<0.05).

*P. tricornutum* cell lines expressing S3-3A or WT showed similar growth under control conditions (Fig. 6A). At day 4, the cultures treated with 10 µM lovastatin did not result in any growth defects in the following 2 days, differently from small-scale pilot experiments (data not shown). Therefore, on day 6 we increased the concentration of lovastatin to 25 µM and we immediately observed a decrease in growth (Fig. 6A) and in photosynthetic efficiency (Fig. S5) on days 7 and 8.

The expression of the ddGFP was pronounced in S3-3A (ddRFPb-AR-2.7 and MBP-3.6-ddGFPa), being slightly lower in DMSO treated cells at day 4 (Fig. 6D). At day 8, the trend was inverted, with the recombinant product being slightly more abundant in DMSO control cultures, where the overall amount of recombinant product substantially increased in both sample sets compared to day 4. In agreement with previous experiments (Fig. 3 and 4), we could not detect GFP signal in WT cultures (Fig. 6A), nor a positive signal in the immunoblot, confirming that the cultures lost the expression of WT (ddRFPb-AR and MBP-ddGFPa) (Fig. 6D). This experiment was carried out 24 months after the above-mentioned screening and localization experiments (Fig. 2 and 5) and it is plausible that transgenic diatoms underwent endogenous silencing mechanisms, or episome instability issues, which are not yet understood but occasionally occur in *P. tricornutum*^75,76^. Nevertheless, we employed this strain as negative controls, as it fulfills the requirements of a conventional ‘empty vector’ control.

The mean fluorescence intensity of S3-3A progressively increased during days 4-6, in correlation with the metabolically active status of cells in this growth phase (Fig. 6A and S5), and the measured relative FPP levels on day 5 and 6 (Fig. 6E), without substantial differences in the presence or absence of lovastatin 10 µM. Cultures expressing S3-3A exhibited higher population heterogeneity (Fig. 6C) plausibly reflecting the different sizes of prenyl phosphate pools in non-synchronized diatom cultures, which include cells of varying ages and metabolic statuses. The biosensor fluorescence reached its maximum on day 6, after which the signal began to decrease (Fig. 6A and C), while the expression of ddRFPb-AR-2.7-MBP-3.6-ddGFPa increased between day 4 and 8 (Fig. 6D) indicating that while the availability of recombinant biosensor was abundant in the cell, the amount of activated conformation declined. This observation appears in contrast with the quantification of prenyl phosphates (GPP and FPP) by LC-MS: while the pools confirm the trend of the biosensor signal on day 5 and 6, on day 8 this increase (Fig. 6E). Interestingly, we could not detect GGPP in any of our samples and GPP only on day 8 (Fig. 6E). On day 8, GPP pools increased to surpass those of FPP, confirming the presence of large pools of GPP in *P. tricornutum*^77^ and a peculiar regulation of the biosynthesis of this compound in late stages of cultivation after treatment with 25 µM lovastatin. In interpreting these results, it is important to consider that the biosensor signal reflected the cytosolic pool of prenyl phosphates, while the metabolites analyzed by LC-MS have been extracted from whole cells and includes the pool contained in the chloroplasts, peroxisomes and possibly other organelles. Hence, discrepancies between these results, such as the one we observed, can be expected. In addition, the drastic decrease in fluorescence signal intensity from day 7 to day 8, despite the increased availability of the recombinant biosensor components (Fig. 6D), could also reflect instability, modification or sequestration of the biosensor component by diatoms in the stationary phase, that may prevent proper assembling and functioning of the recombinant complex.

Interestingly, growth performances in days 7 and 8, and prenyl phosphate pools were both higher in cultures transformed with WT constructs that were no longer functional, compared to those expressing S3-3A, with pools of prenyl phosphates two-fold higher (Fig. 6A and E), suggesting the possibility of direct or indirect effects of the expression of ddRFPb-AR-2.7-MBP-3.6-ddGFPa on diatom physiology and metabolism.

To determine the phenotypic effects of the metabolic perturbation, we measured squalene, the precursor of sterols and direct product of the conversion of cytosolic FPP in *P. tricornutum*, and brassicasterol and campesterol, the main sterols produced by *P. tricornutum*^16,18,71^, as well as the carotenoid pigments fucoxanthin, diatoxanthin, diadinoxanthin and chlorophyll *a* and *c* (Fig. 7) at day 8. We did not observe substantial differences between treated and untreated diatoms in the accumulation of squalene, brassicasterol and campesterol, in cultures expressing WT, while we observed differences between WT and S3-3A cultures in the general response to lovastatin (Fig. 7A and B). While in WT cultures the amount of triterpenoids and carotenoids did not change with the treatment, S3-3A exhibited slightly higher metabolite levels in control conditions, with a significant drop in treated samples (Fig. 7A and B), aligned with typical effects of the inhibition of HMGR^78^. Overall, untreated cultures of cells expressing S3-3A accumulated more total triterpenoids and total carotenoids than those expressing WT. We hypothesize that this observation might be related to the more dynamic fluctuations of prenyl phosphates, compared to those of terpenoid metabolic sinks (i.e. squalene, sterols and carotenoids), which - differently from prenyl phosphates - are not rapidly converted or degraded in the cell. Cells did not divide between days 6 and 8 when treated with lovastatin 25 µM (Fig. 6A), hence their biomass composition did not have sufficient time to change substantially. We hypothesize that in this scenario, cells did not synthesize more triterpenoids on days 7 and 8 and our analysis mostly quantified compounds already synthesized by the cells, until day 6. Interestingly, we observed a similar phenomenon occurring in the biosynthesis of carotenoids, which is fueled by GGPP in the chloroplast^79^, and which we could not detect (Fig. 6E). The inhibition with lovastatin 25 µM resulted in a decrease in the main carotenoids fucoxanthin and diadinoxanthin, which might indicate the occurrence of some compensation effect between the MEP pathway in the chloroplast and the MVA pathway in the cytosol, even if we only observed this effect in diatom cultures expressing S3-3A (Fig. 7B), presumably for the same above-mentioned hypothetical prenyl phosphate sequestration effect by S3-3A. Interestingly, in both diatom cultures, we observed a drastic decrease in chlorophyll *a,* in the presence of lovastatin (Fig. S6B). Finally, it is also possible that cells might encounter reduced availability of prenyl phosphates for further biochemical conversions, while these are bound or ‘trapped’ by the biosensor.

### Considerations, limitations and need for improvements

Our experiments provide an extensive functional investigation and parameterization of S3-3A as an effective biosensing system for the non-invasive, rapid sensing of various prenyl phosphates (GPP, FPP, and GGPP) in *P. tricornutum*. Our results demonstrated its versatility and feasibility for implementing high-throughput screening workflows for strain engineering and addressing fundamental questions about diatom isoprenoid metabolism. To the best of our knowledge, the use of S3-2D and S3-3A biosensors has not been reported in any other organisms *in vivo*, aside from *E. coli*^29^. Hence, it is not known whether their expression could be detrimental to eukaryotic microorganisms. We encountered substantial difficulties working with both S3-2D and S3-3A in diatoms that will require further investigation into the effects of the biosensors in the diatom cellular environment and optimization. We observed overall low conjugation efficiencies compared to those involving other unrelated constructs. Furthermore, cell lines correctly expressing S3-2D and S3-3A often exhibited frequent and drastic growth impairments (data not shown), necessitating the repetition of experiments. In some instances, cells ceased to express WT, S3-2D, or S3-3A, as noted above (Fig. 3, 4 and 6D). These challenges suggest that there may be some interference from the recombinant constructs with diatom physiology. Further investigations involving transcriptomics and metabolomics analyses of the engineered strains may elucidate the factors at play and guide optimization strategies.

### Conclusions

In conclusion, our work represents the first-of-its-kind installation of a protein-based metabolite biosensor capable of detecting various prenyl phosphates in microalgae. We demonstrated that S3-3A is a valid tool for rapid and reliable measurements of the intracellular pool of prenyl phosphate and that it led us to discover novel details in the biosynthesis of isoprenoids in diatoms, such as the involvement of peroxisomes in the MVA pathway. While this exploratory work also highlighted limitations and the need for further investigation and optimization of these biosensors for full deployment in strain engineering workflows, we anticipate that this resource will be of central relevance in the investigation and engineering of the terpenoid metabolism in diatoms.

## Materials and Methods

### Strains and cultivation conditions

*P. tricornutum* strain CCMP632 was obtained from the collection of National Center for Marine Algae and Microbiota (Bigelow Laboratory for Ocean Sciences, USA). Enriched Seawater Artificial Water^80^ (ESAW) liquid medium was used to culture the cells and half-strength ESAW supplemented with 1% agar and 100 μg/ml zeocin (Thermo Fisher Scientific, USA), was used for the selection of transgenic diatoms. In all experiments diatoms were cultivated at 21°C shaking at 95 rpm under 80 μmol photons m^−2^ s^−1^continuous light conditions in a shaking incubator (Innova S44i, Eppendorf, Germany). The volume of cultures was 400 ml in shake flasks, and 200 μl per well in microwell plate assays. The transgenic lines were plated as droplets on solid half-strength ESAW media maintained at 8°C in between the experiments.

### TXTL protein synthesis and *in vitro* assays

TXTL reactions for each plasmid (expressing MBP or AR variants) were prepared separately using the myTXTL linear DNA expression kit (Daciel Arbor Biosciences, USA) according to the manufacturers protocol. The protein concentration was determined by Bradford reagent (Bio-Rad, USA). A master mix was prepared by mixing 1 volume of TXTL extract expressing an MBP variant, 1 volume of TXTL extract expressing an AR variant, and 2 volumes of the PBS/BSA solution. The master mix aliquots of 27 µl were mixed with 3 µl geranyl pyrophosphate (Axon Medchem BV, Netherlands) or farnesyl pyrophosphate (Merck, Germany), each in the concentrations of 0, 0.1, 0.3, 0.5, 1, 2, 5, and 10 µM. The mix was aliquoted as 10 µl in triplicates in a black 384 well-plate (Merck, Germany) and incubated at room temperature for 10 mins. The detailed protocol can be found in Glasgow et al., 2019. A plate reader (BioTek Synergy H1, Agilent, USA) was used to read the ddGFP complementation fluorescence from the top with excitation/emission wavelengths set to 380 nm and 508 nm.

### Cloning and diatom transformation

Plasmids carrying WT, S3-2D and S3-3A were kindly provided by Anum Glasgow (Columbia University, USA), Tanja Kortemme (University of California, San Francisco, USA) and Robert Campbell (University of Alberta, Canada) and the sensor design was as follows: ddRFPb-AR-2.7 and MBP-2.5-ddGFPa for S3-2D, ddRFPb-AR-2.7 and MBP-3.6-ddGFPa for S3-3A and ddRFPb-WTAR and WTMBP-ddGFPa for WT. Primers (Table S2) were designed to include the *u*Loop CD overhangs to the sensor constructs. To remove BsaI site from ddRFPb-WTAR the plasmid was amplified in two fragments using the oligo pairs FP_TxTl9-OP1 RP_TxTl9-OP1 and FP_TxTl9-OP2 RP_TxTl9-OP2 including the overlapping region. The PCR products were used as templates for overlap extension PCR to domesticate ddRFPb-WTAR. All domesticated constructs were cloned as *u*Loop CD L0 parts^20^ for expression in *P. tricornutum*. The coding sequence of *Phatr3_J53939* was ordered from Genewiz (Azenta Life Sciences, USA) with CD overhangs and cloned as L0 *u*Loop part. The assembled L2 plasmids (Table S1) were transformed into *E.coli* Epi300 cells containing conjugative plasmid pTA-MOB^82^ by heat shock, diatoms were transformed by bacterial conjugation method as described previously^83^. After conjugation the transformed diatoms were incubated at 21°C for two days under 80 μmol photons m^−2^ s^−1^ in continuous light conditions. The cells were resuspended in 500 μl ESAW and plated on selection plates containing 100 μg/ml zeocin. Plates were incubated for 15 days at 21°C under 50 μmol photons m^−2^ s^−1^ continuous light conditions until colonies were sufficiently large to be picked. Single colonies were picked and plated in liquid medium in 96-well round bottom plates (Nunclon-treated, Thermo Fisher Scientific, USA) containing 200 μl ESAW with 100 μg/ml zeocin. Independent cell lines were grown for one week and sub-cultured every 4 days to choose the candidates for downstream experiments.

### Flow cytometry analyses

Transgenic diatom cell lines were screened by flow cytometry using Guava EasyCyte flow cytometer system (Guava easyCyte HT, Cytek Biosciences, USA). The GFP fluorescence was measured by the 488-nm blue laser with the Green-B spectral imaging band (525/30 nm). The cells were measured at a flow rate of 0.59 µl/s with 10000/30000 events per sample in triplicates. The data was analyzed in Floreada.

### Exogenous metabolite sensing experiments

Cell lines expressing WT, S3-2D and S3-3A were inoculated in 10 ml of ESAW and grown in 50 ml Erlenmeyer flasks for four days at 21°C in an incubator, shaking at 95 rpm under 80 μmol photons m^−2^ s^−1^ continuous light conditions. On the fourth day, the cultures reached a density of 0.3 OD_750_ and 190 µl culture of each strain was transferred into 96-well plates. 10 µl of the metabolites GPP (Axon Medchem BV, Netherlands), FPP (Merck, Germany), GGPP (Echelon Biosciences, USA) and HMGR inhibitor Lovastatin (Nordic Biosite, Sweden) were added in concentration ranges mentioned in the text and figure legends, and plates were incubated for 3, 6, 24 and 48 hours before measurement by flow cytometry and/or plate reader. The metabolites were dissolved in water and lovastatin in DMSO. GFP fluorescence was measured using a flow cytometer and spectrofluorometrically by plate reader (BioTek Synergy H1, Agilent, USA). The plate reader was used to read the ddGFP complementation fluorescence from the top with excitation/emission wavelengths set to 380 nm and 508 nm.

### Endogenous metabolite sensing experiments

The overall growth performance of WT and S3-3A was monitored over the course of the experiment for 8 days. Cell density was measured spectrophotometrically as OD_750_ and by flow cytometer (BioTek Synergy H1, Agilent, USA) and the photosynthetic activity (quantum yield, Qy) using chlorophyll-*a* pulse amplitude modulated (PAM) fluorometry (AquaPen-C, AP 110-C, Photon System Instruments, Czech Republic). 10 μM of lovastatin (Nordic Biosite, USA) was added on day 4 and the concentration was increased to 25 μM on day 6.

### Intracellular metabolites sampling and extraction

Intracellular metabolites were sampled from day 4 to 8, extracted and analyzed as described previously^84^. The buffer used to extract the samples was 10 µM ampicillin as internal standard in 75% ethanol. The extracted metabolites were snap frozen and stored at −80°C and later concentrated by evaporating under freeze-dryer overnight. The samples were resuspended in 100 μl 50% MeOH and analyzed by LC-MS.

### LC-MS analysis

The prenyl phosphate pools were analyzed using liquid chromatography (LC) coupled to ion mobility (IM) quadrupole time of flight (qTOF) mass spectrometry (MS) as described previously^85^. The instrumentation consisted of an Agilent 1290 Infinity II series UHPLC system hyphenated with an Agilent 6560 IM-qTOF with a Dual Agilent Jet Stream Electron Ionization source. LC separation was performed on an InfinityLab Poroshell 120 HILIC-Z, 2.1 mm × 50 mm, 2.7 μm UHPLC column (Agilent Technologies 689775–924) coupled to an InfinityLab Poroshell 120 HILIC-Z, 3.0 mm × 2.7 μm UHPLC guard column (Agilent Technologies 823750–948, USA). A 3.5 min gradient was run using organic buffer (acetonitrile) combined with an aqueous buffer with high pH (10 mM ammonium acetate, pH 9) for negative ionization mode. Data was acquired using MassHunter Data Acquisition 10.0 software on 1 μl of sample separated on the column with a flow rate of 800 μl min^-1^. Data were acquired in the 50 to 1700 m/z range, with an MS acquisition rate of 0.8 scans/s. The raw data files were processed using the Agilent MassHunter software suite. Features were annotated based on accurate mass and CCS values using a manually curated reference library consisting of GPP, FPP and GGPP. Peak areas were calculated using the MassHunter Qualitative Analysis 10.0 software.

### Confocal microscopy analyses

Diatom cells expressing WT, S3-3A and overexpression of PtMVK (Phatr3_J53929) tagged with mVenus at an OD_750_ of 0.2 were used for confocal microscopy analyses. The samples were imaged with AX Confocal/Multiphoton system (Nikon, Japan) using plan Apo λ 100x oil-immersion objective with 1.49 NA (Nikon, Japan) and at a resolution of 1024 × 512 pixels. ddGFP or mVenus were excited by 488 nm argon laser and chlorophyll autofluorescence with 561 nm laser and the fluorescence emission of ddGFP or mVenus was detected at a bandwidth of 500-520 nm and of chlorophyll at 625–720 nm. Light was passed through the transmitted light channel. The images were processed by ImageJ software.

### Determination of PTS2 in Phatr3_J53939 (PtMVK)

The coding sequence of Phatr3_J53939 obtained from the *Ensembl* database (https://protists.ensembl.org/Phaeodactylum_tricornutum/Info/Index), was searched manually for the nonapeptide consensus sequence [RKSH]-[LVIQAT]-X_5_-[HDQ]-[LAF] within the first 50 amino acids. The consensus sequence was obtained from various known PTS2-like peptides from *Arabidopsis thaliana*, *Saccharomyces cerevisiae* and *Homo sapiens*^66,67^.

### Protein extraction, SDS-PAGE and immunoblotting

Two ml diatom cultures from day 4 and 8 treated with and without lovastatin were harvested by centrifugation at 3500g at 4°C for 10 mins, the pellet was washed once with 2 ml PBS, snap frozen in liquid nitrogen and stored at −80°C until protein extraction. For extraction of crude protein the protocol was modified from^86^, the pellets from biological triplicates were pooled, amounting to 8 samples in total, and thawed in 500 μl lysis buffer (50 mM Tris-HCl pH 7.5, 150 mM NaCl, 1 mM EDTA, 0.1% (*v*/v) Triton X-100, protease inhibitor cocktail tablet (Thermo Fisher Scientific, USA)) and lysed by sonication (Branson Digital Sonifier, USA) for 3 min on ice with 30 secs ON/OFF. The sonicated cells were centrifuged at 20,000*g* and 4°C for 2 hours and proteins in the supernatant was precipitated with 100% acetone overnight and the dried pellets were dissolved in 2.5 M urea. The protein concentration of the samples was determined by Bradford protein assay kit (Bio-Rad, USA) according to the manufacturer’s guidelines. The crude proteins were denatured by addition of SDS-PAGE Laemmli sample loading buffer (1:4) (Bio-Rad, USA) containing β-mercaptoethanol (1:10) for 10 min at 95°C. Proteins were separated on 4–15% SDS-PAGE (Mini-PROTEAN® TGX Stain-Free™ pre-cast gels, Bio-Rad, USA) at 200 V for 30 mins in Tris/glycine/SDS buffer (Bio-Rad, USA). The proteins were transferred on to precut PVDF membrane (Trans-Blot Turbo Midi 0.2 µm PVDF Transfer Packs, Bio-Rad, USA) using a Trans-Blot® Turbo™ Transfer system (Bio-Rad, USA). The membrane was soaked in MilliQ water for 2 mins and then in 100% ethanol for 30 secs and completely dried. The membranes were blocked for 2 hours in 5% *w*/*v* skim milk in PBS + 0.1% Tween 20 (Merck, Germany) (PBST) buffer. After blocking, the membrane was washed with TBST buffer (1x Tris buffered saline (Bio-Rad, USA) + 0.05 % Tween 20 (Merck, Germany)) and probed with an anti-GFP tag monoclonal antibody (Thermo Fisher Scientific, USA) at a dilution of 1:1000 overnight at 4°C and washed in TBST. Secondary detection was performed using a goat anti-Mouse IgG, IgM-HRP (Thermo Fisher Scientific, USA) at a dilution of 1:5000 followed by Clarity ECL Western Blotting substrate (Bio-Rad, USA) detection using a ChemiDoc MP (Bio-Rad, USA) imaging system.

### Triterpenoid extraction and quantification by GC-FID

50 ml diatom cell cultures of 0.8 OD_750nm_ were harvested by centrifugation for 10 minutes at 3500g and 21°C. The pellet was washed once in PBS buffer and flash frozen in liquid nitrogen and stored at −80°C until use. The frozen pellets were freeze-dried overnight in a freeze dryer (laboratory freeze dryer Alpha 1-2 LSC basic, Martin Christ, Germany) and the weights of the dried pellets were recorded. The freeze-dried cells were lysed by incubation at 95°C for 10 minutes in 250 μl each of 40% KOH and 50% ethanol. Then, 900 µl of hexane was added and the organic layer was transferred to amber vials and the extraction was repeated and the extracts were combined. The extracted solvent was evaporated under nitrogen air. The samples were derivatized at 70°C for one hour with 20 µl of pyridine and 100 µl of N-methyl-N-(trimethylsilyl)-trifluoroacetamide. The samples were analyzed by an Agilent 7890B GC with flame ionization detector with a DB-5 column (30.0 m x 0.25 mm x 0.25 µm) using Helium as carrier gas at a constant flow of 1ml/min. The GC-FID was set up as follows: oven temperature post injection was set at 80°C for a minute and ramped to 280°C at 20°C/min, held at 280°C for 45 min, ramped to 320°C at 20°C/min, held at 320°C for a min and cooled to 80°C at 50°C/min at the end of the run. Triterpenoid peaks were identified based on the retention time of the standard triterpenoids. Peak areas were normalized to dry weight of the samples.

### Photosynthetic pigments extraction and analysis

50 ml diatom cultures were harvested, washed and stored as above. The frozen cells were freeze-dried overnight, and the weights of the dried pellets were recorded. The dried pellets were resuspended in 95% cold methanol with 2% ammonium acetate and the extracts were transferred to amber vials and incubated in dark at −20°C for one hour. The supernatant was extracted by centrifugation at 6000g and filtered through 0.2 µm PTFE 13 mm filter to remove the cell debris before measuring at the HPLC and stored at −80°C until analysis. The pigment absorbance spectra were determined at 400 nm using an Agilent HPLC system with a YMC HPLC column 150 x 3.0 mm (S-3 µm). The column temperature was maintained at 40°C with a flow rate of 0.5 ml/min. The two eluents were A: acetone/methanol 40/60 (v/v) and B: water/acetone 40/60 (v/v) with elution gradient as follows: 0-22 min from 0 to 30% B, 22-41 min from 30 to 10 % B, and 45-55 min from 10 to 60% B. Pigments were identified based on the retention times of the standards and UV absorbance spectra at 400 nm. The peak area was normalized to dry weight of the samples.

### Statistical analyses and reproducibility

All data was analyzed and generated in Prism (GraphPad). Error bars represent the standard deviation of the mean. The significance evaluated by one-way ANOVA. The number of replicates is indicated in the figure legends.

## Author Contribution

MF conceived this project; PP and MF designed the experiments; PP carried out all the experiments with contributions from MF. PP analyzed the data with contributions from MF. TM developed the LC-MS methods and performed the intracellular prenyl phosphate quantification analyses, reviewed and edited the manuscript. PP and MF wrote the manuscript. All authors read and approved the manuscript.

## Acknowledgements

This work was supported by VILLUM FONDEN (grant number 3721 to MF) and by the SDU Climate Cluster (SCC) through a Research Infrastructure Grant to MF. We would like to thank Anum Glasgow (Columbia University, USA), Tanja Kortemme (University of California, San Francisco, USA) and Robert Campbell (University of Alberta, Canada) for kindly providing the biosensor constructs. We would also like to thank Aniruddh Murali (PhyLife, SDU) for helping with data analysis and Marion Sébire for her help.

## Abbreviations

MVA: mevalonate
MEP: methyl erythritol
GPP: geranyl diphosphate
FPP: farnesyl diphosphate
GGPP: geranyl geranyl diphosphate
HMGR: 3-hyroxy-3-methyl-glutaryl-CoA
GFP: green fluorescent protein
WT: wild-type

