## Supplementary Material File 1 for "Real-time tracking of intracellular prenyl phosphate pools in the marine diatom *Phaeodactylum tricornutum* with a metabolite protein-based biosensor"

**Supporting information**

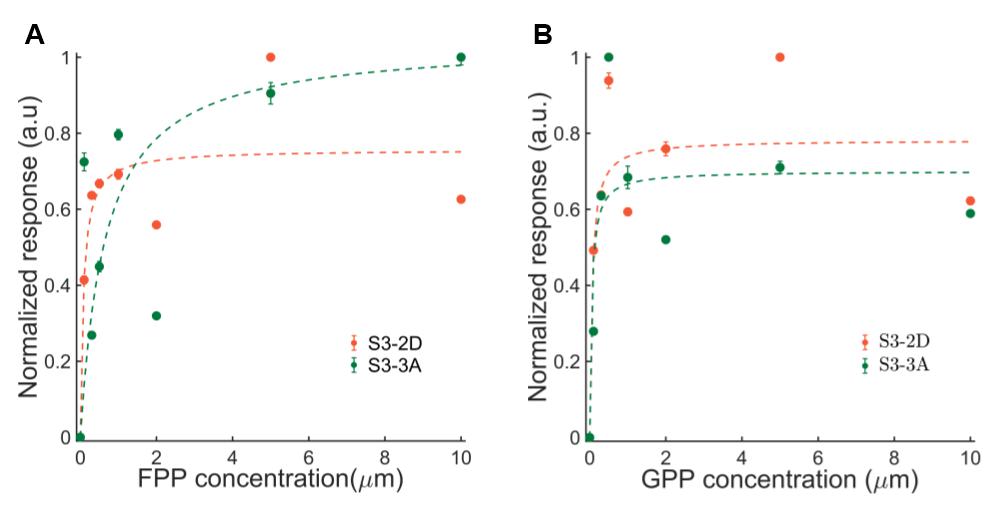

**Figure S1**. Dose-response curve of cell-free extracts of biosensor variants after incubation for 10 mins with different concentrations of (A) FPP and (B) GPP.

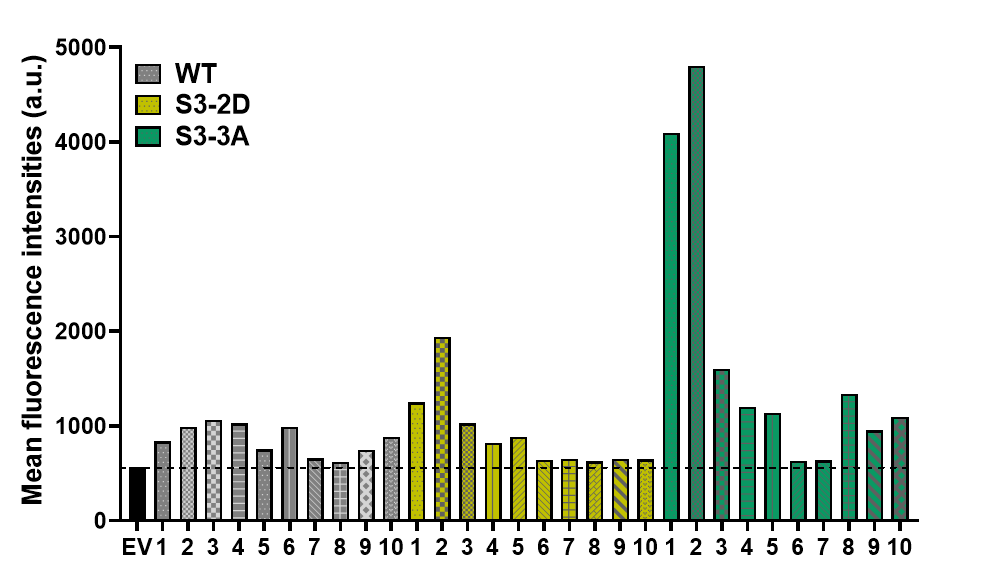

**Figure S2.** Flow cytometry-based screening of 30 transgenic diatom cell lines expressing WT, S3-3A or S3-2D biosensors and EV control. The bars represent the mean fluorescence intensity of 10,000 cells for each sample, numbers on the x-axis indicate independent cell lines.

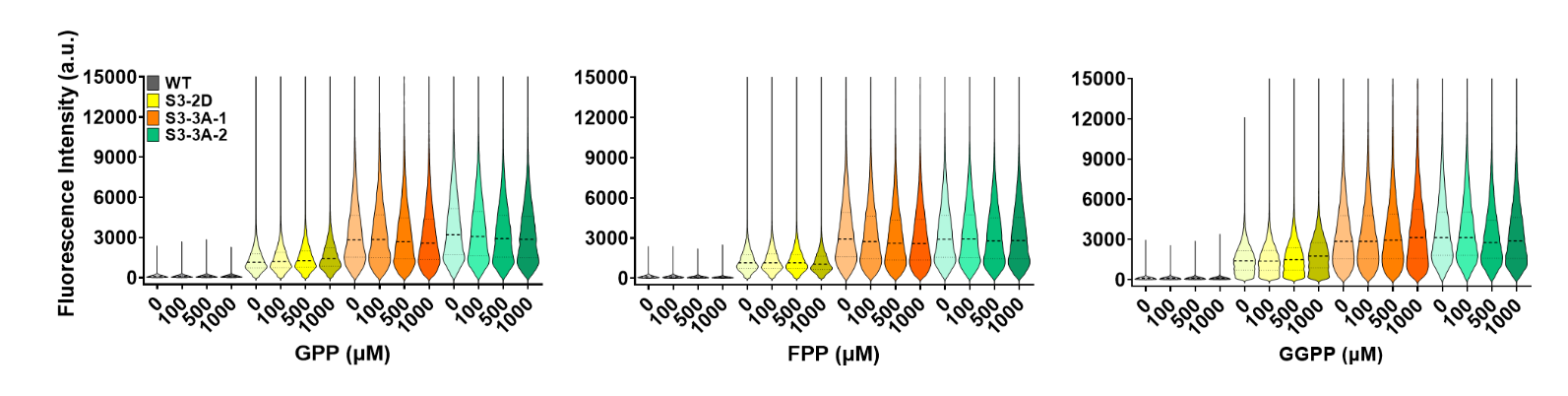

**Figure S3.** Violin plots depicting the activity of WT, S3-2D and S3-3A after 6 hours of incubation after the addition of macro-concentrations of GPP, FPP and GGPP (n=30,000). Dashed black lines represent median and grey dotted lines represent upper and lower quartiles.

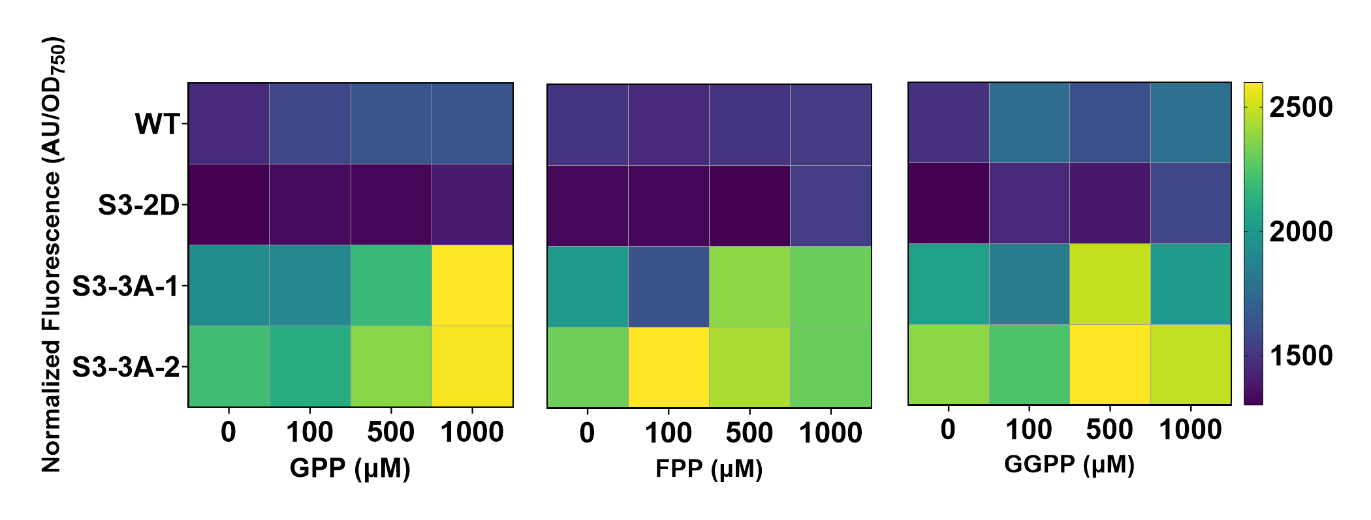

**Figure S4.** Mean fluorescence intensity normalized by OD^750^ of diatom cell lines expressing either S3-2D, S3-3A or WT dimerization complexes, in the presence of GPP, FPP and GGPP at different concentrations (0-1000 µM), detected by a spectrofluorometer after 24 hours of incubation (n=3).

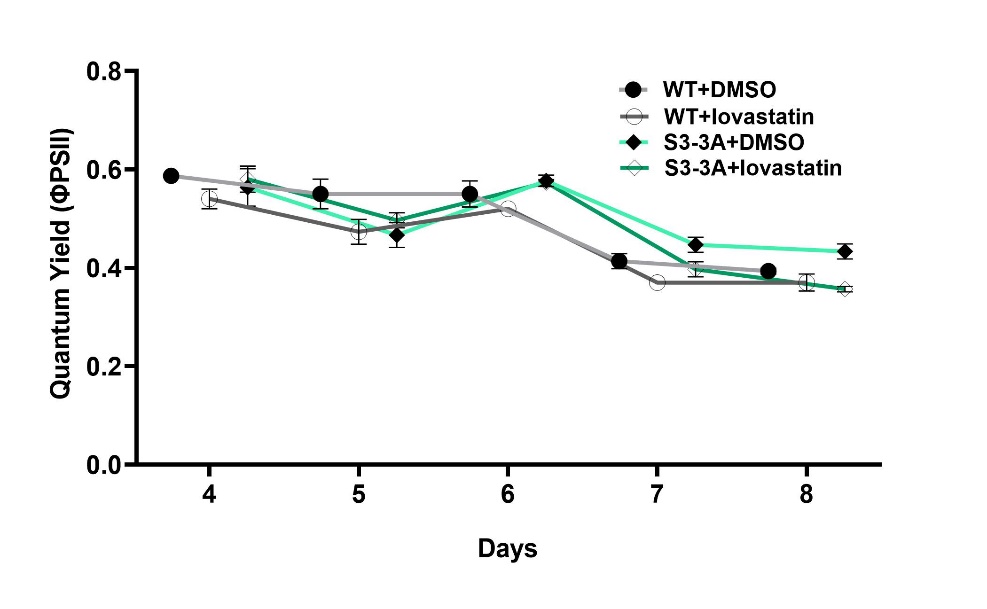

**Figure S5**. Effective quantum yield of transgenic diatom strains expressing either WT or S3-3A, either treated with lovastatin 10µM (days 4-6), 25 µM (days 6-8) or DMSO (solvent control) (n=3).

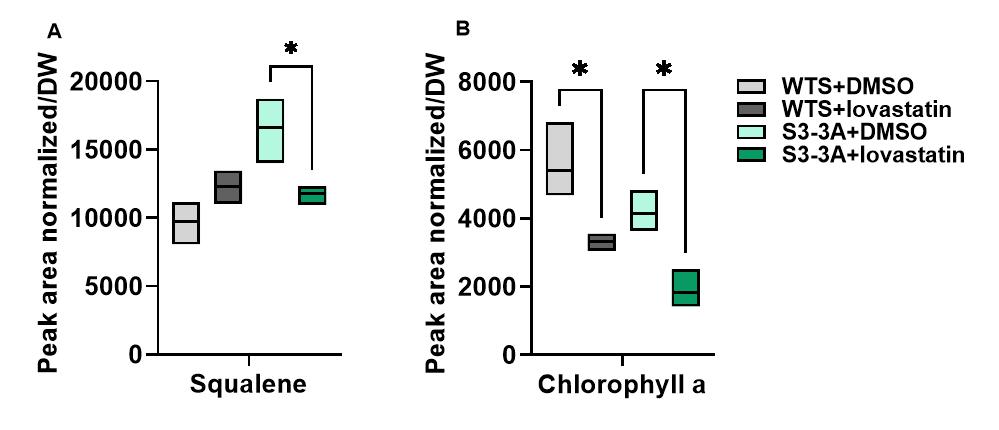

**Figure S6**. Relative quantification of (A) Squalene and (B) chlorophyll *a* in transgenic diatom strains expressing either WT or S3-3A, either treated with lovastatin 10µM (days 4-6), 25 µM (days 6-8) or DMSO (solvent control). Extraction and quantification of squalene and pigments were carried out on day 8 (n=3).

**PHATR3_J53929**

MAGTPLINSRSVCASAPGKAILFGEHAVVYGEPAVAAALDDLRIFVLFTPTSHNSTHASVIRVVMPDLPTPVDFALPVSLFTSMESTNTPPTPDDATVLAQLLHTADPTLDDFSVQALTPVLYLCNQILLKPLRFRSESNPSADGGYELWVRSQDLPVGAGLGSSAAFGVACAAALIQYRQNLTATHAGTAAGTTSTNTPIQYGPPDSATLSEIDRYAYYSEILLHGTPSGIDNAVSAHGGAIIFTKDVASTNGTVKMEHLTPSDDLTLSLVYTHVPRSTKTLVAGVRHFYQRHVGFVTLILEAMGAIARDFEQAIRDRTNAHHGDHASDSHDFGERVLTMVRTNQYLLQAVGVSHPSLDHVCAVVHEHFRDYGAAKLTGAGGGGCAFVLWKPGLAADVLATQRQRLQYALTTTLTPSPPYRYQCLASVVGGEGVLFLPAGDFPWDATTPSSATPSRSATNVGRLVSALTVSALVATTWMVWRSIGTKAR

**Figure S7**. Amino acid sequence of the predicted mevalonate kinase (Phatr3_J53929), with the predicted PTS2 highlighted in red

**Supplementary table 1**: List of uLoop assembly parts constructed in this study.

| **No** | **Level** | **Position** | **Name** | **Description** |
| --- | --- | --- | --- | --- |
| 1 | L0 | CD | ddRFPb-AR-2.7 | S3-2D and S3-3A |
| 2 | L0 | CD | MBP-2.5-ddGFPa | S3-2D |
| 3 | L0 | CD | MBP-3.6-ddGFPa | S3-3A |
| 4 | L0 | CD | ddRFPb-WTAR | WT |
| 5 | L0 | CD | WTMBP-ddGFPa | WT |
| 6 | L1 | pCAo-2 | pCAoL1-2_p49202-ddRFPb-AR-2.7-FcBPt | S3-2D and S3-3A |
| 7 | L1 | pCAo-3 | pCAoL1-3_p49202-MBP-2.5-ddGFPa-FcBPt | S3-2D |
| 8 | L1 | pCAo-3 | pCAoL1-3_p49202-MBP-3.6-ddGFPa-FcBPt | S3-3A |
| 9 | L1 | pCAo-2 | pCAoL1-2_p49202-ddRFPb-WTAR-FcBPt | WT |
| 10 | L1 | pCAo-3 | pCAoL1-3_p49202-WTMBP-ddGFPa-FcBPt | WT |
| 11 | L2 | pCAe-1 | pCAe-1_pCAoL1-1-PTCv2-pCAoL1-2-p49202-ddRFPb-AR-2.7-FcBPt-pCAoL1-3_p49202-MBP-2.5-ddGFPa- FcBPt-pCAoL1-4-Spacer4 | S3-2D |
| 12 | L2 | pCAe-1 | pCAe-1_pCAoL1-1-PTCv2-pCAoL1-2-p49202-ddRFPb-AR-2.7-FcBPt-pCAoL1-3_p49202-MBP-3.6-ddGFPa- FcBPt-pCAoL1-4-Spacer4 | S3-3A |
| 13 | L2 | pCAe-1 | pCAe-1_pCAoL1-1-PTCv2-pCAoL1-2-p49202-ddRFPb-WTAR-FcBPt-pCAoL1-3_p49202-WTMBP-ddGFPa- FcBPt-pCAoL1-4-Spacer4 | WT |

**Supplementary table 2**: Primers used in this study.

| **No** | **Name** | **Primer** |
| --- | --- | --- |
| 1 | FP_TxTL4/6/10_uloop-CD | AAGCTCTTCATCCAATGTCTAAAATCGAAGAAGGTAAACTGGTAATC |
| 2 | RP_TxTL4/6/10_uloop-CD | TTGCTCTTCTTCGACCTGAGCTGCCTGTGCTACCTGTAC |
| 3 | FP_TxTL3/9_uloop-CD | AAGCTCTTCATCCAATGGTGAGCAAGGGCGAGGAAACC |
| 4 | RP_TxTL3/9_uloop-CD | TTGCTCTTCTTCGACCTGAGTTCAGTTTCTGCAGGATTTCAGCC |
| 5 | RP_TxTL3_uloop-CD | TTGCTCTTCTTCGACCTGACTAGGCCGCTTTCTGCAGCAC |
| 6 | FP_TxTL3/9-OP1 | GTGAGCAAGGGCGAGGAAACCATCAAAGAG |
| 7 | RP_TxTL3/9-OP1 | GCCATGTAGATGGTTTCGAACTCCACCAGGT |
| 8 | FP_TxTL3/9-OP2 | GAAACCATCTACATGGCCAAGAAGCCCGTG |
| 9 | RP_TxTL3/9-OP2 | GTTCAGTTTCTGCAGGATTTCAGCCAGGTC |
| 10 | RP_TxTL3-OP2 | CTAGGCCGCTTTCTGCAGCACTTCAGC |
